# Exogenous ECM in an environmentally-mediated *in vitro* model for cardiac fibrosis

**DOI:** 10.1101/2024.08.20.608840

**Authors:** Natalie Pachter, Kristen Allen, Tracy A Hookway

**Author notes:** **Corresponding Author:** Tracy A. Hookway, PhD, 65 Murray Hill Road, Vestal, NY 13850.

## Abstract

Few clinical solutions exist for cardiac fibrosis, creating the need for a tunable *in vitro* model to better understand fibrotic disease mechanisms and screen potential therapeutic compounds. Here, we combined cardiomyocytes, cardiac fibroblasts, and exogenous extracellular matrix (ECM) proteins to create an environmentally-mediated *in vitro* cardiac fibrosis model. Cells and ECM were combined into 2 types of cardiac tissues-aggregates and tissue rings. The addition of collagen I had a drastic negative impact on aggregate formation, but ring formation was not as drastically affected. In both tissue types, collagen and other ECM did not severely affect contractile function. Histological analysis showed direct incorporation of collagen into tissues, indicating that we can directly modulate the cells’ ECM environment. This modulation affects tissue formation and distribution of cells, indicating that this model provides a useful platform for understanding how cells respond to changes in their extracellular environment and for potential therapeutic screening.

## Introduction

The cardiac environment broadly consists of 2 components- cells and extracellular matrix (ECM). Cardiac cells include cardiomyocytes (CMs), the contractile cells of the heart, and cardiac fibroblasts (CFs), which are responsible for the production and turnover of ECM. Other cardiac cell types include endothelial cells, epicardial cells, neurons, macrophages, pericytes, stromal cells, and smooth muscle cells (1,2). The ECM surrounds cardiac cells, providing structural support, promoting cellular communication, and providing a dynamic environment of growth factors, proteases, cytokines, and chemokines (1,3). The ECM consists of both a non-fibrillar basement membrane portion and a dense interstitial matrix. The basement membrane, containing fibronectin, laminin, collagen IV, and proteoglycans, forms an interface between CMs and the interstitial matrix and holds growth factors and cytokines until their release is signaled (3,4). The interstitial matrix consists of the fibrillar component of the cardiac ECM and contains mostly collagens I and III, which provide mechanical support (3,4). While collagen I makes up around 85% of the cardiac ECM and is also the main component of deposited ECM during fibrosis (1,5), non-fibrillar proteins within the basement membrane have been shown to contribute to myocardial remodeling after injury, meaning they also serve an important role in cardiac fibrosis (4).

Cardiac fibrosis is characterized by increased deposition of extracellular matrix (ECM) proteins in the heart, resulting in stiff, nonfunctional scar tissue and impaired cardiac contraction (6). Cardiac fibrosis may result from an acute ischemic injury, such as myocardial infarction, but can also develop as a consequence of other conditions, including diabetes, cardiomyopathy, and hypertension (7,8). Excess ECM deposition is driven by cardiac fibroblasts (CFs). In the healthy heart, in addition to ECM maintenance, CFs also provide structural support and assist with cellular crosstalk and propagation of electrical signal throughout the heart (5). When stressed, CFs transition to myofibroblasts, which are highly migratory and produce excess disorganized ECM, largely collagen I (5).

Few treatments exist for cardiac fibrosis due to the heart’s limited regenerative capacity and long-term cardiac toxicity of approaches such as inhibition of transforming growth factor β (TGF-β) (6,7). Additionally, the dynamic ECM deposition and remodeling that occurs after injury is crucial for stabilization of the injury site, meaning this process cannot be fully inhibited (9). Further, the importance of collagen in both the heart and elsewhere in the body complicates the use of collagen degradation as an appropriate treatment without careful targeting and control (10). This creates a need for a reliable *in vitro* model that may be used to develop new treatments for cardiac fibrosis.

While animal models allow for assessment of interactions between organ systems and observations over longer timescales, significant differences between humans and other animals often result in poor clinical trial outcomes. Upwards of 50% of therapeutics that show efficacy in animal models do not translate this efficacy to humans, and close to 95% of drugs fail in human clinical trials (11,12). *in vitro* models, while limited in representing organism-level interactions, allow for the use of human cells, higher throughput, and greater tunability of cell composition and environment. Further, differentiation of induced pluripotent stem cells (iPSCs) to cardiomyocytes (CMs) provides an on-demand cell source not subject to the availability of rare and transient primary cardiac tissue. Genetic editing of iPSCs also allows for simple visualization of cardiac proteins such as cardiac troponin I (TNNI) (13) or calcium flux activity (14) in differentiated cells. iPSCs may also enable modeling with patient-specific cell lines for highly relevant efficacy screening. Further, the availability of iPSC lines powers the study of a variety of cardiac conditions which feature dynamic extracellular environments, such as cardiac fibrosis.

To date, most *in vitro* cardiac fibrosis models lack either multiple types of human cardiac cells (15–18), a 3D environment enabling more *in vivo*-like cell interactions (17–19), or a defined ECM environment (20). Most existing models also indirectly induce fibrosis using a chemical mediator, usually TGF-β, which signals fibroblasts to transition to myofibroblasts and thereby deposit more ECM (16,17,21). Given the importance of TGF-β signaling in fibrosis, this indirect induction is a valuable approach to examine fibrosis *in vitro*, but lacks control over which specific ECM proteins are present, their ratios, and their amounts. Further, providing cells with ECM outside of endogenous proteins produced by CFs may be important for understanding cell behavior, given the *in vivo* proximity and communication of cardiac cells and ECM. The field lacks an environmentally-mediated fibrosis model which investigates the effects of tuned ECM environments on cardiac cells.

### Significance

Here, we present a 3D *in vitro* cardiac fibrosis model combining hiPSC-derived CMs and primary human cardiac fibroblasts with exogenously added ECM proteins in ratios that mimic the human *in vivo* environment. We generated 2 different types and scales of 3D cardiac tissues-aggregates and rings. Aggregates, requiring a lower cell number, allow for higher throughput screening of ECM conditions while rings feature mechanical support via a central post, greater opportunity for cellular alignment, and a larger tissue that promotes more complex cellular interactions.

We show that exogenous ECM modulates the structure of these tissues. At the smaller tissue scale, collagen has a severe negative impact on tissue formation, but does not significantly impact contractile function. In larger tissues, collagen had a less drastic impact on formation, but a similarly insignificant impact on contractile function. Overall, we show that exogenous ECM proteins can modulate 3D cardiac tissues and provide a tunable model to investigate individual contributors to cardiac fibrosis, including specific proteins and their ratios. This environmentally-mediated model enables examination of how cells respond to changes in their ECM environment as opposed to a chemical inducer of cardiac fibrosis.

## Methods

### Cardiomyocyte differentiation

Human iPSCs were maintained prior to differentiation on Geltrex or Matrigel-coated tissue culture plates, fed daily with Essential 8 medium (Gibco), and passaged every 3 days to maintain pluripotency. iPSCs genetically encoded with a GCaMP-6f calcium reporter (Gladstone Institutes) or with a cardiac troponin I (TNNI) reporter (Allen Institute) were differentiated to CMs via Wnt pathway modulation (22,23). For GCaMP iPSCs, differentiation was initiated on Day 0 at 95-100% confluence with 12 μM GSK inhibitor CHIR99021 (Selleckchem) for exactly 24 hours. 5 μM Wnt inhibitor IWP2 (Selleckchem) was given on Day 3 for 48 hours. Base media of RPMI/B27 (Corning, Gibco) without insulin was used until Day 7, then replaced with RPMI/B27 containing insulin and changed every 3 days. CMs began spontaneous beating on Day 7-10. For TNNI iPSCs, differentiation was initiated on Day 0 at 80-90% confluence with 7.5 μM CHIR99021 for exactly 48 hours, followed by 7.5 μM IWP2 on Day 2 for 48 hours. RPMI/B27-was replaced with RPMI/B27 containing insulin on Day 6, then changed every 3 days. CMs began spontaneous beating on Day 12-15. CMs used in cardiac tissues were between Day 23-38.

### Cardiac fibroblast culture

Primary human adult CFs (Cell Applications) were maintained in 2D on tissue culture-treated plates prior to use in 3D cardiac tissues. CFs were fed every 3 days with fibroblast growth media from Cell Applications or in-house CF media (DMEM/F12, 10% fetal bovine serum, 1% NEAA, 1% L-glutamine, 1% penicillin-streptomycin) and passaged at ∼90% confluence. CFs used in cardiac tissues were between passages 4-6.

### Cardiac aggregate formation

GCaMP or TNNI CMs and CFs were combined (3:1) with collagen I (bovine or rat tail, Corning), laminin (mouse, Corning), collagen and laminin (2:1, mimicking the ratio of these proteins *in vivo* (24)), or Geltrex (Gibco) at 0.1, 0.25, and 0.5 mg/mL. Cell-ECM mixtures were seeded into round bottom, ultra low attachment 96 well plates (Costar) at 10,000 cells per 20 μL volume (Fig. 1). Cell-only aggregates with no exogenous ECM were used as a control. The plate was gently centrifuged after seeding, and cells were allowed to form aggregates for 24 hours before 200 μL media (RPMI/B27 with insulin and 1% penicillin-streptomycin) was gently added to the side of the wells. Formation was assessed 24 hours after seeding via bright field microscopy (Nikon Eclipse Ti2, Andor Zyla camera, NIS Elements software) and aggregates were periodically imaged for 1 week after seeding. For all aggregate experiments, n ≥ 5 for each condition with at least 3 independent experiments.

**Figure 1:**
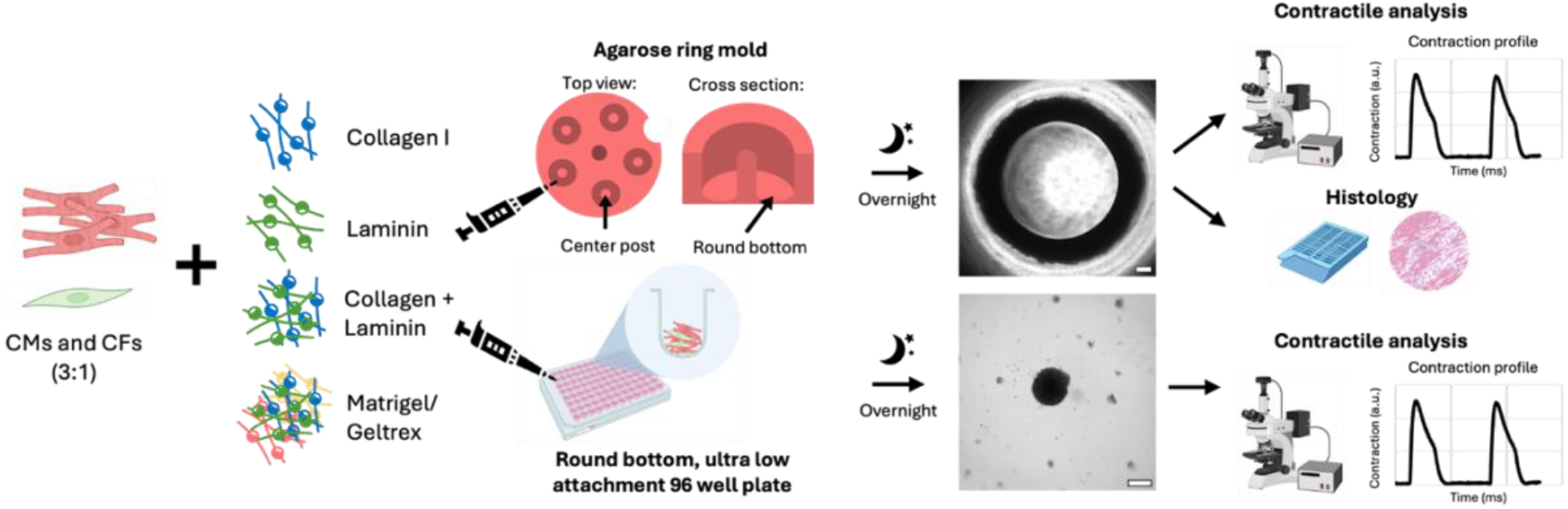
Schematic of cardiac ring and aggregate formation. Scale bars = 250 μm.

### Cardiac tissue ring formation

GCaMP CMs and CFs were combined (3:1) with collagen I (rat tail), laminin (mouse), collagen and laminin in equal parts, collagen and laminin (2:1, mimicking the ratio of these proteins *in vivo* (24)), or Matrigel (Corning) at 0.1 and 0.25 mg/mL. Cell-ECM mixtures were seeded into custom agarose ring molds at 70 μL and 900,000 cells per ring and allowed to self-assemble into rings overnight (25) (Fig. 1). Formation was assessed 24 hours after seeding via bright field microscopy and rings were imaged again at 2, 4, and 8 days after seeding. n = 5 for each condition.

### Contractile analysis

24-48 hours following recovery of spontaneous beating, high frame rate (67-100 fps) calcium flux or bright field videos were captured for contractile analysis of GCaMP and TNNI-tagged aggregates and rings, respectively (Nikon Eclipse Ti2, Andor Zyla camera, NIS Elements software). For each aggregate, a single, square (100-200 μm^2^) uniformly beating region of interest (ROI) was captured. For each ring, 4 uniformly beating square (400 μm^2^) ROIs were captured. Videos were converted to .avi format in Fiji, then analyzed with the MuscleMotion Fiji plugin (26). Contraction profiles generated by MuscleMotion were manually verified to ensure that all beats were properly captured. MuscleMotion parameters (speed window, estimated peak width, and Gaussian blur) were optimized and applied consistently within a dataset to generate the greatest possible number of properly analyzed videos. Quantitative contractile data were then statistically analyzed in RStudio.

### Statistical analysis

Analyzed contractile parameters included contraction duration, time-to-peak, relaxation time, contraction amplitude, and peak-to-peak time. For aggregates, n ≥ 5 for each condition with 3 independent experiments. For rings, n = 5 for each condition. For cardiac tissue rings with multiple captured ROIs, all ROIs were averaged to generate a single data point per ring. Normality and homogeneity were assessed with a Shapiro-Wilk test and Bartlett test, respectively, with p > 0.05 indicating normality or homogeneity. Normally distributed and homogeneous datasets were analyzed with a one-way ANOVA and Tukey post hoc test while non-normally distributed and/or non-homogeneous datasets were analyzed with a Kruskal-Wallis test with Benjamini-Hochberg adjustment and Dunn post hoc test. p < 0.05 was considered statistically significant.

### Whole mount staining and confocal microscopy

Three days after seeding, aggregates were fixed in suspension in 4% paraformaldehyde for 45 minutes, then washed 3 times in PBS, permeabilized in 0.2% Triton X 100 for 15 minutes, washed again in PBS, and blocked for 1 hour in 1% bovine serum albumin. Primary antibody (cTnT, Abcam, 1:200) was added and incubated overnight at 4°C, then washed. Secondary antibody (Alexa Fluor 594, Invitrogen, 1:500) was added and incubated overnight at 4°C protected from light, then washed. Phalloidin 488 (Invitrogen, 1:100) and Hoechst (Invitrogen, 1:2000) were incubated for 1 hour at room temperature, protected from light. Aggregates were placed on a rocker for all incubation steps. Aggregates were washed and mounted on glass slides with Prolong Gold antifade reagent (Invitrogen). Once dry, aggregates were imaged on a Zeiss LSM880 confocal microscope.

### Histology

7-10 days after seeding, cardiac tissue rings were fixed in 10% formalin for 1 hour, then removed from their agarose molds (27) and embedding in paraffin. 5 μm sections (Thermo Scientific HM 325) at a 5° angle were collected on glass slides. Slides were stained with hematoxylin (5 min) & eosin (1 min) or picrosirius red (0.1 wt. % Direct Red 80 and Fast Green in picric acid, 30 min), then analyzed via bright field imaging (Nikon Tn2 Eclipse).

## Results

### Collagen negatively impacted aggregate formation

Aggregates containing CMs, CFs, and exogenous ECM were seeded in ultra-low attachment, round bottom 96 well plates and allowed to form overnight before initial formation was assessed with phase microscopy. Successful aggregate formation was defined as one or several dense, compact cell structures with no web-like projections. Aggregates that formed as flat, non-compact cell regions, with web-like projections, or as cells scattered throughout the well were considered not successfully formed. Successfully formed aggregates had an average diameter of 300 μm.

Collagen negatively impacted cardiac aggregate formation (Fig. 2). Poor compaction was observed 24 hours after seeding in aggregates containing collagen, whether alone or together with laminin (Fig. 2a, c). In these poorly compacted aggregates, cells flattened against the surface of the ultra-low attachment well into “patches” instead of forming dense structures (Fig. 2d, 0.25 and 0.5 mg/mL collagen). This occurred across a range of collagen concentrations (0.1, 0.25, and 0.5 mg/mL) (Fig. 2d). Some aggregates also formed as small clumps of cells scattered throughout the well which did not compact (Supplemental video 2). 2 days after seeding, a small percentage (6.5%) of collagen containing aggregates developed a “webbed” phenotype, where some cells formed a compact center but other cells and ECM components protruded outward from the center in a spiderweb-like structure, adhering to the side of the ultra-low attachment well (Fig. 2d, 0.1 mg/mL collagen). Collagen also affected the spatial organization of aggregates. Those without collagen exhibited dense cellular cores and those with collagen exhibited more diffuse outgrowths (Fig. 2).

**Figure 2:**
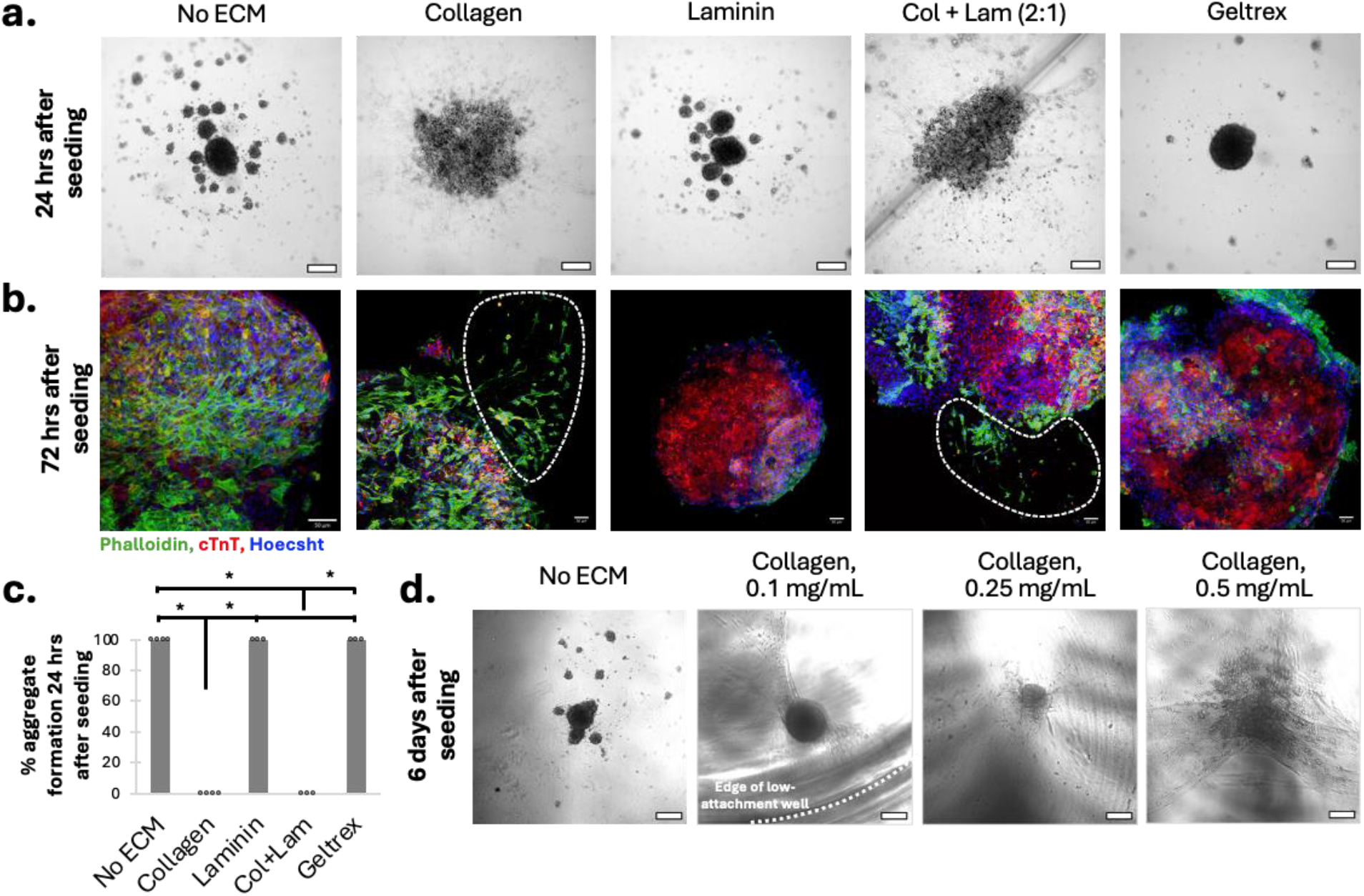
Collagen impairs aggregate formation. **(a)** Phase imaging of aggregates 24 hours after seeding (scale bars = 250 μm) and **(b)** confocal imaging of aggregates 72 hours after seeding (scale bars = 50 μm). All ECM groups 0.25 mg/mL. Regions outlined with white dashed lines indicate diffuse outgrowths lacking cTnT signal. **(c)** Percent aggregate formation by ECM type at 0.25 mg/mL, 24 hours after seeding. Circles indicate independent experiments, each with n ≥ 5. *p < 0.05. **(d)** Phase imaging of aggregates 6 days after seeding, showing abnormal formation at additional collagen I concentrations (scale bars = 250 μm).

### Collagen modulates ring formation less drastically than in aggregates

24 hours after seeding, ring formation was consistent across all ECM conditions (Fig. 3a). All rings had an average outer diameter of approximately 3 mm and were contracted around the center agarose post. After 4 days in culture, changes in ring remodeling were visible in all groups with uneven ring thickness emerging in some groups by day 4 of culture. Webbing was visible in collagen-containing rings, and in rings without collagen, cells begin to separate into 2 “phases” with less defined edges (Fig. 3c).

**Figure 3:**
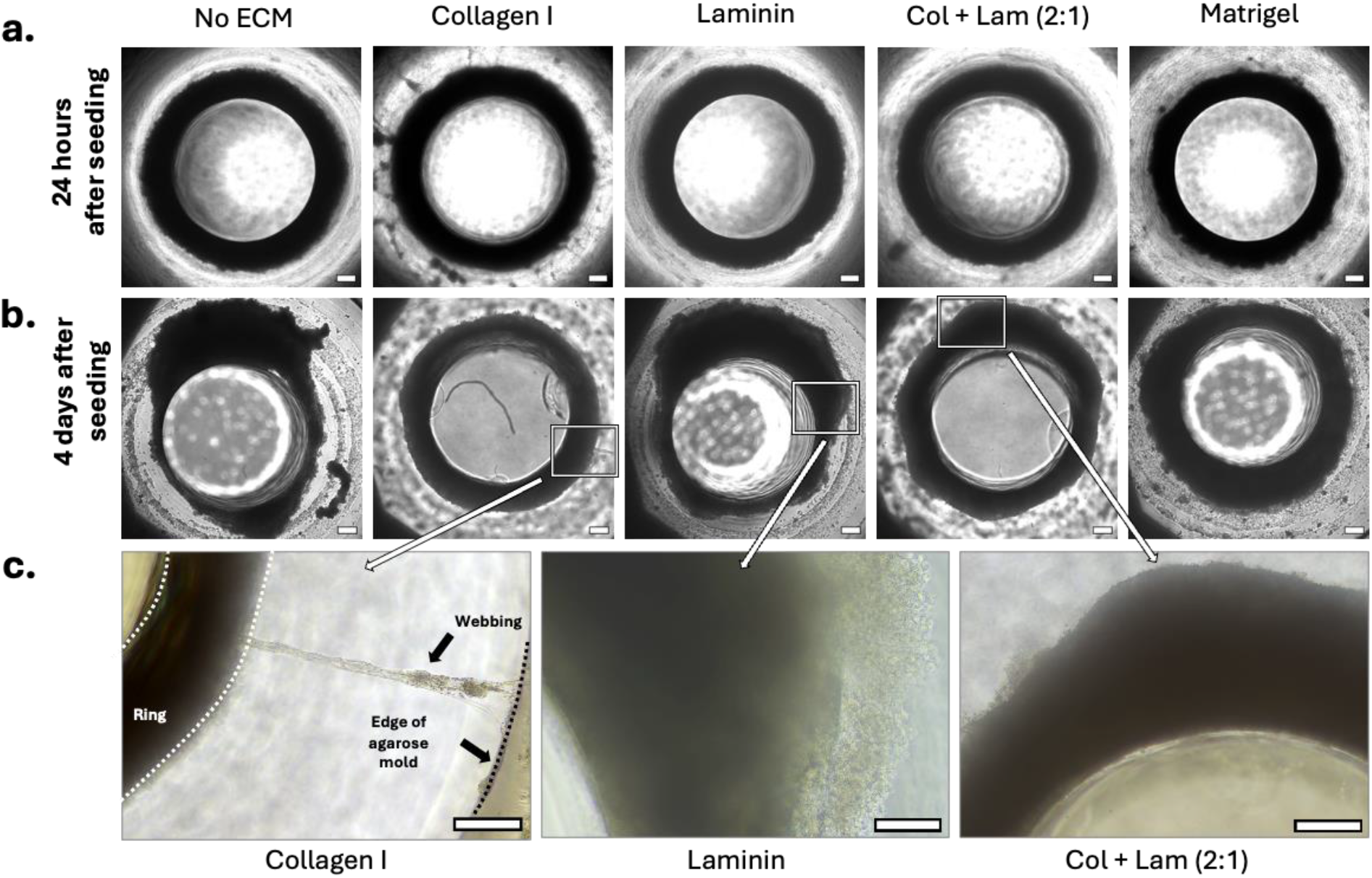
Collagen changes how rings remodel over 4 days. Ring formation **(a)** 24 hours after seeding and **(b)** 4 days after seeding. **(c)** Enlarged sections of images in (b) to show webbing caused by collagen and differences in spatial organization between rings with and without collagen. Scale bars = 250 μm. All ECM added at 0.25 mg/mL.

### Collagen does not impair aggregate contractile function

Although tissue formation was variable across different ECM conditions, it was also important to investigate contractile properties of cardiac aggregates to assess their function. Notably, even poorly formed aggregates, including webbed, flattened patch, and scattered cell morphologies, exhibited robust beating (Fig. 4b). Collagen did not impair contraction amplitude, i.e. beat strength was unchanged (Fig. 4c). Collagen also did not affect contraction duration or time-to-peak. In aggregates containing both collagen and laminin, relaxation time was decreased as compared to collagen alone, laminin alone, or rings with no exogenous ECM (Fig. 4d).

**Figure 4:**
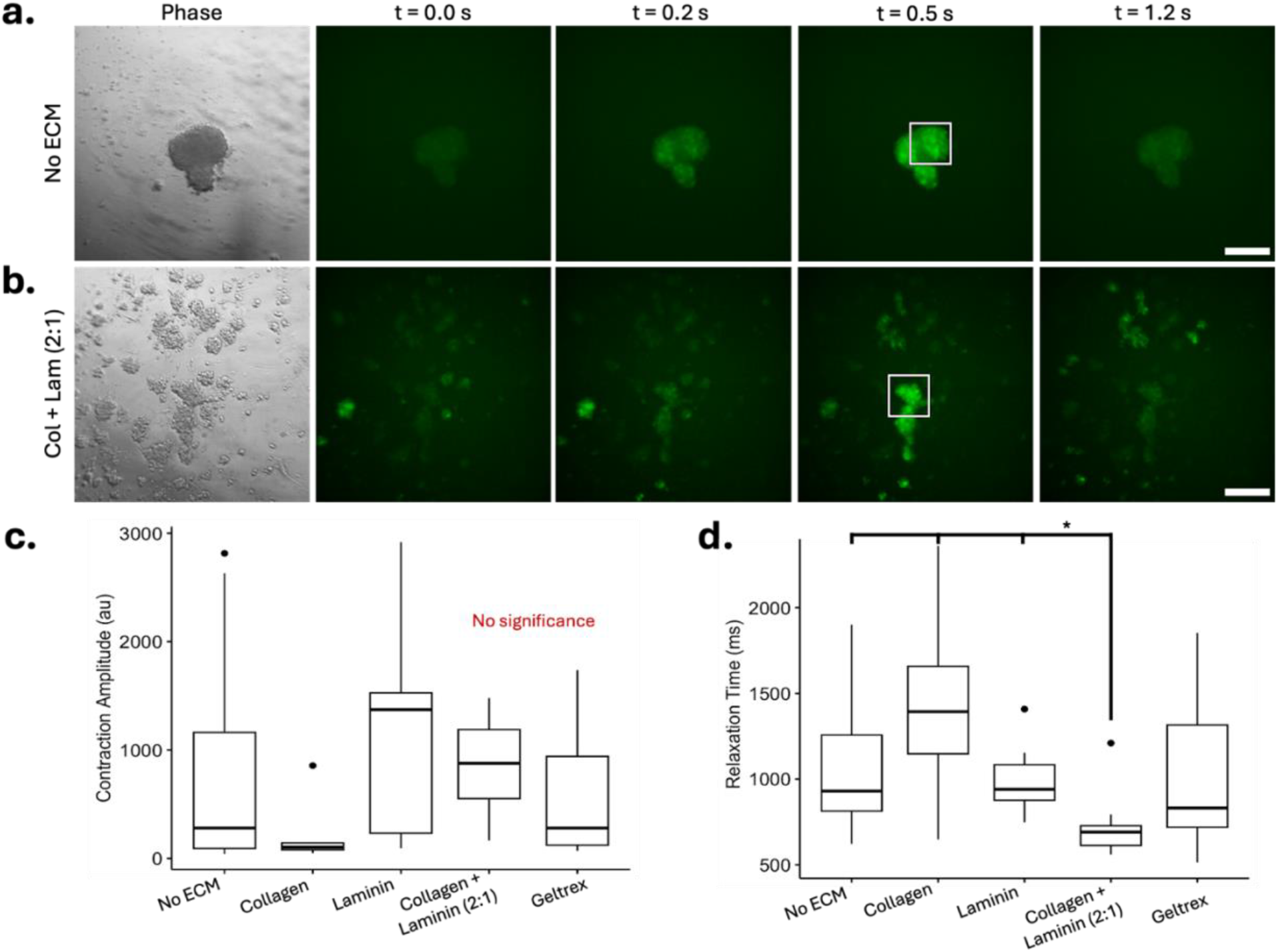
Collagen did not impair beating in aggregates. Calcium flux timelapses and corresponding phase images of aggregates (72 hours after seeding) containing **(a)** no ECM and **(b)** 0.25 mg/mL collagen I + laminin (2:1). White frame indicates representative uniformly beating region of interest that was analyzed with MuscleMotion to provide contractile data. Scale bars = 250 μm. **(c)** Contraction amplitude and **(d)** relaxation time data from aggregates containing 0.25 mg/mL ECM. No significant differences noted in contraction amplitude, but aggregates containing collagen and laminin had decreased relaxation time compared to no ECM aggregates or aggregates with collagen or laminin alone. *p < 0.05.

### Collagen does not drastically impair ring contractile function

Similarly, contractile analysis of the larger, ring-shaped tissues was performed to understand the functional consequences of ECM incorporation. Qualitatively, rings of all ECM conditions exhibited robust beating with periodic flux. Quantitatively, rings containing collagen exhibited decreased contraction amplitude with respect to rings containing only laminin or no exogenous ECM. Rings containing collagen also exhibited significantly shorter relaxation times as compared to most other ECM groups, including rings with no exogenous ECM. Videos of beating tissues are shown in supplementary data (Supplemental videos 1-4).

### Histology shows incorporation of collagen into rings

In order to take a closer look at the interior organization of the tissues, histological sections were analyzed for overall tissue structure. High cellularity was observed in cardiac tissue rings (Fig. 6a). In rings containing collagen, however, clear acellular regions were visible, usually localized to one side of the ring (Fig. 6a). In the collagen containing groups, picrosirius red staining indicated that the areas of lower cellularity contain high levels of collagen, likely representing the added exogenous matrix (Fig. 6b). Of note, significant diffuse collagen was observed in rings containing laminin, Matrigel, or no exogenous ECM.

**Figure 5:**
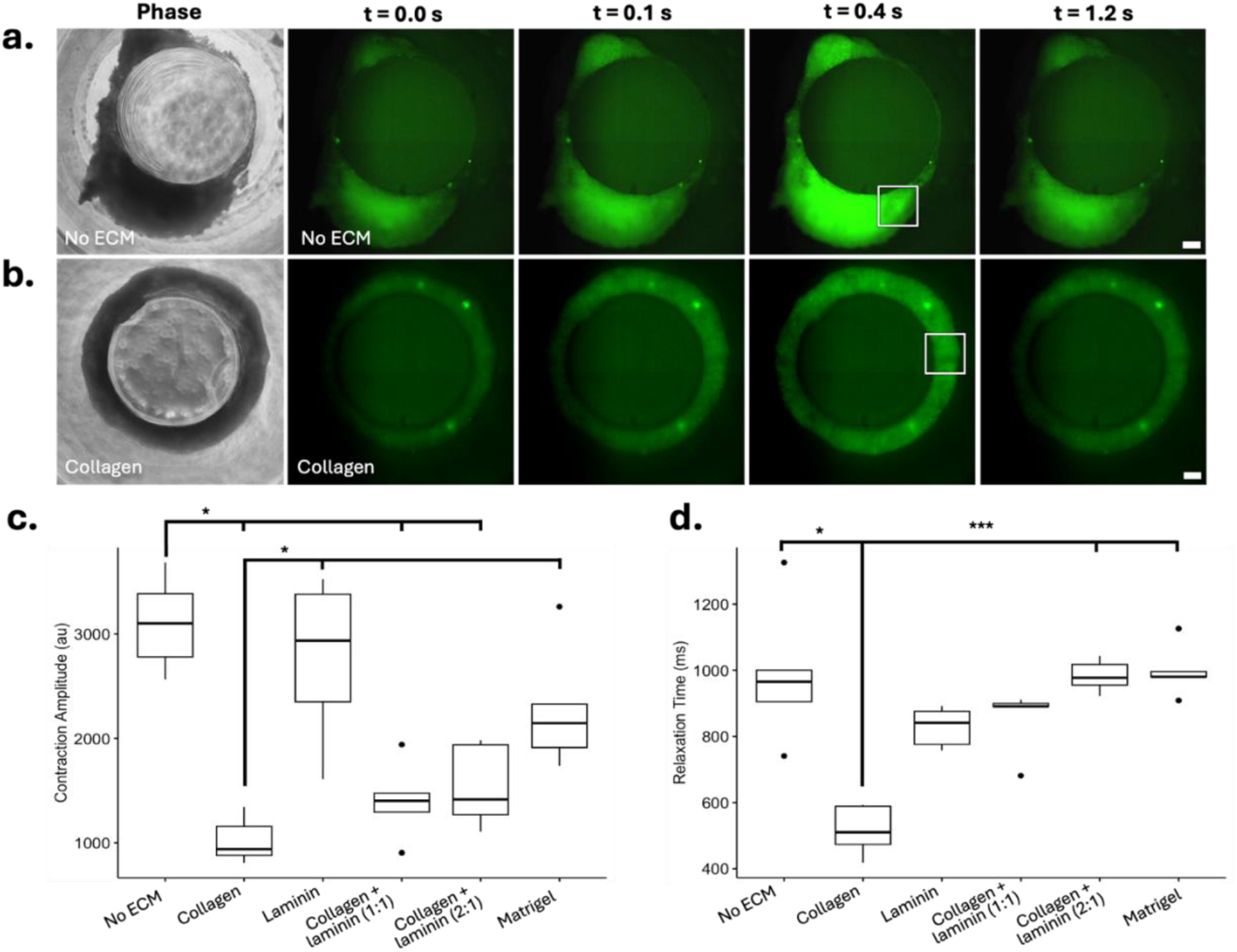
Collagen did not drastically impair beating in rings. Calcium flux timelapses and corresponding phase images of rings (8 days after seeding) containing **(a)** no ECM and **(b)** 0.25 mg/mL collagen I + laminin (2:1). White frame indicates representative uniformly beating region of interest that was analyzed with MuscleMotion to provide contractile data. Scale bars = 250 μm. **(c)** Contraction amplitude and **(d)** relaxation time data from aggregates containing 0.25 mg/mL ECM. All collagen-containing groups exhibited decreased contraction amplitude with respect to rings containing no ECM or laminin alone. However, rings containing collagen also exhibited decreased relaxation time than several other groups, including rings with no added ECM. *p < 0.05, ***p < 0.005.

**Figure 6:**
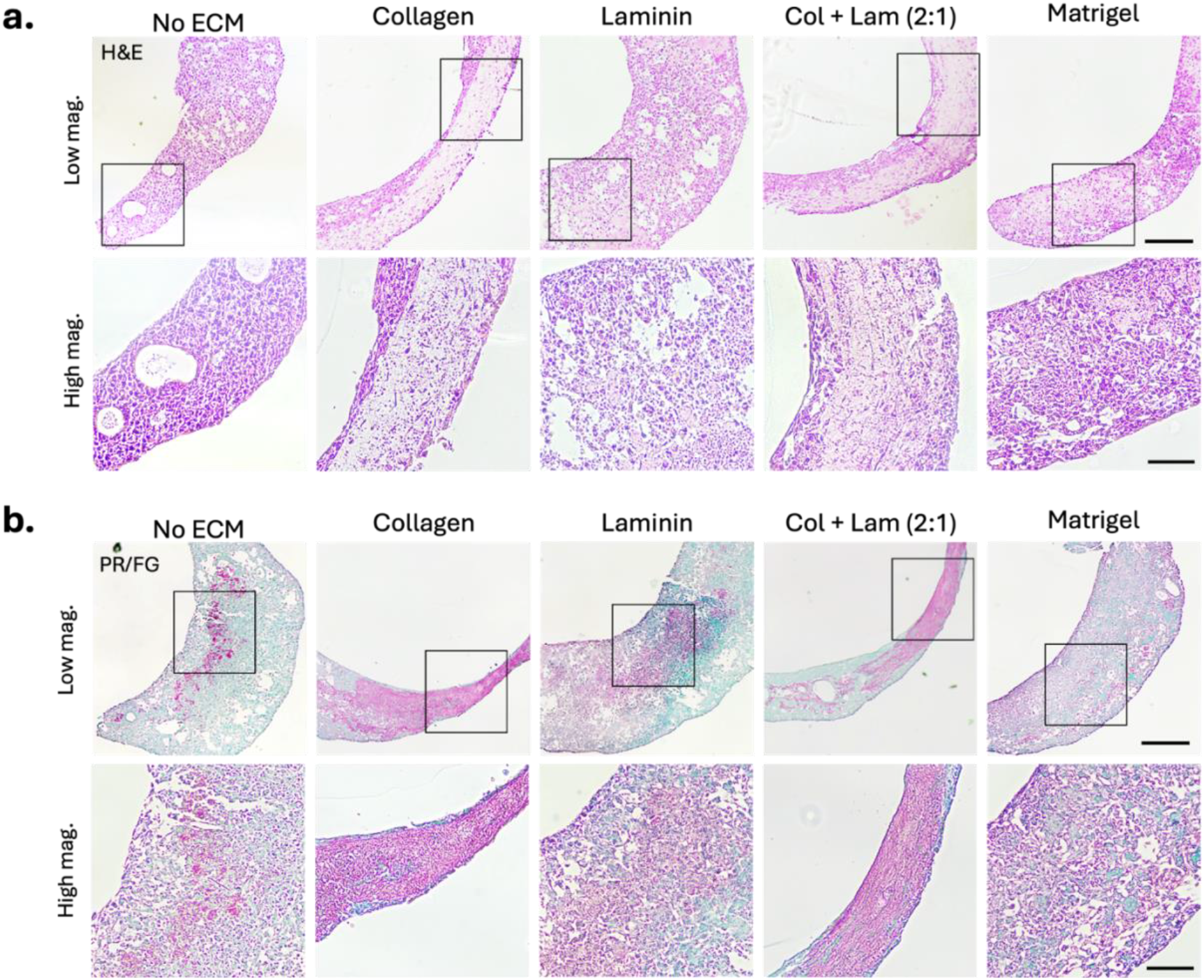
Histological analysis of cardiac tissue rings. **(a)** Hematoxylin & eosin staining of cardiac tissue rings containing 0.25 mg/mL ECM. Hematoxylin (nuclei) is shown in purple and eosin (nonspecific extracellular proteins) is shown in pink. **(b)** picrosirius red & fast green staining of rings containing 0.25 mg/mL ECM. Fast green (nuclei) is shown in green and direct red 80 (collagen) is shown in pink/red. Black boxes indicate areas of low magnification images that correspond to higher magnification field of view. Low magnification scale bars = 250 μm. High magnification scale bars = 100 μm.

## Discussion

Collagen is used broadly in many cardiac tissue engineering applications, including as a component of tissue engineered scaffolds and as a substrate for direct cell seeding (15,28–30). However, here we see variable cell responses to the presence of collagen as compared with other types of ECM. In aggregates, collagen prevented cells from compacting into dense tissues within the first 24 hours of formation (Fig. 2a). This non-compaction occurred whether collagen was added alone or together with laminin. Additional parameters were also modified to ensure that this poor formation was due to collagen, including using a different lot of collagen, multiple iPSC lines, and changing aggregate seeding method and volume, but this poor formation in the presence of collagen persisted (Supplemental Fig. 1). This may be analogous to other studies reported in literature wherein CMs seeded in collagen matrices exhibited improved tissue compaction when cell density was increased and collagen concentration was decreased (15,31).

Several days after seeding, collagen-containing aggregates presented more spatially disorganized morphologies (Fig. 2b, d). These included webbing, wherein a small number of cells compacted in the center, but additional cells and ECM proteins protruded from the center and attached to the side of the low-attachment well, and patches, wherein cells spread out on the surface of the low-attachment well instead of compacting into an aggregate. Immunostaining and confocal microscopy suggest that the webbed areas and peripheries of patches are not primarily composed of CMs and instead composed of CFs and ECM proteins, as exhibited by the lack of cTnT staining in these areas (Fig. 2b, indicated by white dashed regions). This spreading and/or CF migration may be an analogous response to fibroblasts spreading within collagen gels in hydrogel-based models. Other studies have noted spreading of dermal fibroblasts when embedded in collagen gels as well as spreading of foreskin fibroblasts in nested collagen matrices (32,33).

Here, we assessed tissue formation across both millimeter (rings) and micron (aggregates) scales. In rings, formation was uniform across all ECM conditions 24 hours after seeding (Fig. 3a). By day 4 after seeding, cells began to reorganize and ring shape became more inconsistent in rings without collagen (Fig. 3b). Cells also began to separate into 2 “phases” in rings without collagen (Fig. 3c). At this timepoint, collagen-containing rings exhibited webbing, though to a lesser extent than aggregates. By day 4, bright field imaging showed strands of cells and ECM protruding from the main ring and adhering to the side of the agarose ring mold (Fig. 3c). Reorganization of cells and collagen is also reported in other studies using analogous ring platforms (27,34).

In both tissue types, aggregates and rings, collagen impacted formation. Interestingly, collagen caused adhesion to traditionally non-adhesive surfaces, ultra-low attachment wells and agarose ring molds, both of which are designed to discourage adhesion. This may indicate that collagen is providing a substrate for cell and/or protein attachment.

Collagen had a more drastic negative effect on formation of aggregates as compared to rings. 24 hours after seeding, 100% successful aggregate formation was seen in control aggregates with no exogenous ECM as compared to 0% successful formation in collagen-containing aggregates (Fig. 2c). However, rings of all ECM conditions had nearly 100% successful ring formation. Even rings that later developed webbing were largely uniform (Fig 3). This difference in how collagen affected the formation of these 2 tissue types may be attributed to the relative size of the tissues. This is supported by literature reporting increased tissue compaction of cardiac cells in collagen scaffolds when cell density was increased (31). Aggregates, containing 10,000 cells each, were seeded in a volume of 20 μL while rings, containing 900,000 cells each, were seeded in a volume of 70 μL. Exogenous ECM concentrations were calculated in mg/mL based on these respective volumes, meaning that the amount of exogenous ECM relative to cell number was markedly greater in aggregates than in rings (500 cells for every 1 μL ECM in aggregates vs. 13,000 cells for every 1 μL in rings). Even if CFs are producing some endogenous collagen, as suggested by picrosirius red staining of rings without exogenous ECM (Fig. 6b), the relatively small number of cells in an aggregate means that exogenous ECM still constitutes a much greater fraction of total ECM in the aggregate system as compared to rings. Higher concentrations of ECM may be needed to achieve similarly drastic webbing and patch phenotypes in rings. This is supported by preliminary studies using 0.5 mg/mL collagen in rings, which resulted in inconsistent ring thickness (Supplemental Fig. 2). Similar studies using collagen typically use it in scaffolds at higher concentrations for increased structural integrity. These concentrations typically range from 0.5 - 6 mg/mL (31,35–37).

In addition to tissue formation, assessment of the contractility of our tissues in the presence of exogenous ECM is important to the understanding their functional behavior. This understanding of contractile function will help improve relevance of this model to *in vivo* function of the fibrotic heart. CMs take up and release calcium as they contract through excitation-contraction coupling (38). The use of GCaMP-tagged iPSC-CMs allows for fluorescent imaging of calcium handling and quantitative analysis of this contraction. More mature and robustly contractile CMs are expected to show decreased time-to-peak, relaxation time, and contraction duration as well as increased contraction amplitude, i.e. stronger and faster individual beats (38).

Despite collagen negatively affecting aggregate formation, even poorly formed, non-compact tissues still exhibited robust beating (Fig. 4b). No significant differences in contraction amplitude were observed between any ECM groups, indicating that beat strength was not affected. This correlates with other 3D engineered heart tissue models discussed in literature, wherein increasing the total amount of collagen in the model did not change contractile force of the tissues (15).

Aggregates containing both collagen and laminin had shorter relaxation times compared to most other ECM groups (Fig. 4d). This was a preliminary indication that multiple ECM types at a physiologically relevant ratio may be beneficial to some parameters of contractile function. This finding motivated the inclusion of an additional condition in subsequent ring experiments wherein collagen and laminin were included both in equal parts and in a physiologically relevant ratio (2:1).

In rings, contraction amplitude was decreased in those containing collagen (Fig. 5c), but collagen did not completely impair calcium flux activity (Fig. 5, Supplemental Video 4). Interestingly, rings containing collagen had decreased relaxation time as compared to most other ECM groups, indicating an improvement in this beating parameter (Fig. 5d). Additional studies are required to determine whether collagen and laminin in a physiologically relevant ratio is beneficial for contractile function at additional tissue scales. Ultimately, collagen and other ECM types did not drastically affect contractile function despite their influence on tissue structure.

Here, we have shown that inclusion of exogenous ECM proteins in *in vitro* cardiac tissues can modulate their formation while maintaining contractile function. This environmentally-mediated *in vitro* model is a useful addition to existing models that use TGF-β to chemically induce fibrosis (16,17,21). While additional experiments are needed to directly compare these 2 types of models in depth, we have observed that the formation of tissues in response to the addition of collagen differs greatly from formation in the presence of TGF-β (Supplemental Fig. 3). While aggregates containing collagen did not compact, those with TGF-β exhibited normal compaction. Those with both collagen and TGF-β exhibited similar formation to that of aggregates containing collagen alone. This indicates that environmental mediation and chemical mediation have different effects on cell behavior, and inclusion of both types of stimuli may be very important in accurately representing fibrosis *in vitro*.

Because stiffness is a key element in fibrosis, future studies will focus on determining whether exogenous ECM increases the stiffness of these tissues. Crosslinking is another important factor in stiffness; covalent bonds, or crosslinks, between collagen fibrils increase their tensile strength and make them more resistant to degradation (39). Some clinical and *in vitro* evidence suggests that the degree of collagen crosslinking may be more directly correlated to tissue stiffness than absolute collagen amount (40,41). Further studies may include chemical mediation of crosslinking and subsequent stiffness measurements. Additional optimizations that would improve the relevance of this model include the use of additional cardiac cell types such as immune and stromal cells, more complex contractile analysis using electrical stimulation, and optimized ratios of ECM that mimic healthy and fibrotic environments.

In this study, we have presented a tunable *in vitro* model for cardiac fibrosis that enables targeted analysis of specific ECM types as well as their ratios and amounts. Aggregates provide a simplified, higher-throughput platform for testing a wide range of ECM conditions without requiring extremely high cell numbers. Cardiac tissue rings offer a larger and more complex *in vitro* tissue with a central post providing mechanical support for cells and greater opportunity for cellular alignment. Both rings and aggregates allow customization of the ECM environment and may also be used to look at individual cell types and ratios. With further optimization, this model will help the field’s understanding of fibrotic mechanisms and provide a platform for higher throughput drug screening, accelerating therapeutic development.

## Acknowledgments

The authors would like to thank Dr. Bruce Conklin for providing the GCaMP6f-WTC11 iPSCs. The authors would also like to thank the Binghamton University Health Sciences Core Facility, Haleema Qamar for assistance with confocal microscopy, and Jacob Gershon for pilot studies with tissue rings. Figure 1 schematic was created in BioRender.

## Author Contributions and Consent Statement

TAH and NP designed the study, interpreted results, and prepared the manuscript. NP performed experiments and NP and KA performed data analysis. All authors consent to publication of this article.

## Data availability

Original datasets may be provided upon request from the authors.

## References

1. Pagliarosi O, Picchio V, Chimenti I, Messina E, Gaetani R. Building an Artificial Cardiac Microenvironment: A Focus on the Extracellular Matrix. Front Cell Dev Biol [Internet]. 2020 Sep 4 [cited 2020 Oct 6];8. Available from: https://www.ncbi.nlm.nih.gov/pmc/articles/PMC7500153/

2. Picchio V, Floris E, Derevyanchuk Y, Cozzolino C, Messina E, Pagano F, et al. Multicellular 3D Models for the Study of Cardiac Fibrosis. Int J Mol Sci. 2022 Oct 1;23(19):11642.

3. Silva AC, Pereira C, Fonseca ACRG, Pinto-do-Ó P, Nascimento DS. Bearing My Heart: The Role of Extracellular Matrix on Cardiac Development, Homeostasis, and Injury Response. Front Cell Dev Biol. 2021;8:1705.

4. Chute M, Aujla P, Jana S, Kassiri Z. The Non-Fibrillar Side of Fibrosis: Contribution of the Basement Membrane, Proteoglycans, and Glycoproteins to Myocardial Fibrosis. J Cardiovasc Dev Dis. 2019 Sep 23;6(4):35.

5. Hinderer S, Schenke-Layland K. Cardiac fibrosis – A short review of causes and therapeutic strategies. Adv Drug Deliv Rev. 2019 Jun 1;146:77–82.

6. Raziyeva K, Kim Y, Zharkinbekov Z, Temirkhanova K, Saparov A. Novel Therapies for the Treatment of Cardiac Fibrosis Following Myocardial Infarction. Biomedicines. 2022 Sep 2;10(9):2178.

7. Garoffolo G, Pesce M. From dissection of fibrotic pathways to assessment of drug interactions to reduce cardiac fibrosis and heart failure. Curr Res Pharmacol Drug Discov [Internet]. 2021 [cited 2021 Dec 19];2. Available from: http://www.ncbi.nlm.nih.gov/labs/pmc/articles/PMC8663973/

8. Sharma A, De Blasio M, Ritchie R. Current challenges in the treatment of cardiac fibrosis: Recent insights into the sex-specific differences of glucose-lowering therapies on the diabetic heart: IUPHAR Review 33. Br J Pharmacol [Internet]. [cited 2022 May 18];n/a(n/a). Available from: https://onlinelibrary.wiley.com/doi/abs/10.1111/bph.15820

9. Scridon A, Balan AI. Targeting Myocardial Fibrosis-A Magic Pill in Cardiovascular Medicine? Pharmaceutics. 2022 Jul 30;14(8):1599.

10. McKleroy W, Lee TH, Atabai K. Always cleave up your mess: targeting collagen degradation to treat tissue fibrosis. Am J Physiol-Lung Cell Mol Physiol. 2013 Jun;304(11):L709–21.

11. Van Norman GA. Limitations of Animal Studies for Predicting Toxicity in Clinical Trials: Is it Time to Rethink Our Current Approach? JACC Basic Transl Sci. 2019 Nov 1;4(7):845–54.

12. Seyhan AA. Lost in translation: the valley of death across preclinical and clinical divide – identification of problems and overcoming obstacles. Transl Med Commun. 2019 Nov 18;4(1):18.

13. Grancharova T, Gerbin KA, Rosenberg AB, Roco CM, Arakaki JE, DeLizo CM, et al. A comprehensive analysis of gene expression changes in a high replicate and open-source dataset of differentiating hiPSC-derived cardiomyocytes. Sci Rep. 2021 Aug 4;11(1):15845.

14. Mandegar MA, Huebsch N, Frolov EB, Shin E, Truong A, Olvera MP, et al. CRISPR Interference Efficiently Induces Specific and Reversible Gene Silencing in Human iPSCs. Cell Stem Cell. 2016 Apr 7;18(4):541–53.

15. van Spreeuwel ACC, Bax NAM, van Nierop BJ, Aartsma-Rus A, Goumans MJTH, Bouten CVC. Mimicking Cardiac Fibrosis in a Dish: Fibroblast Density Rather than Collagen Density Weakens Cardiomyocyte Function. J Cardiovasc Transl Res. 2017 Apr;10(2):116–27.

16. Mainardi A, Carminati F, Ugolini GS, Occhetta P, Isu G, Diaz DR, et al. A dynamic microscale mid-throughput fibrosis model to investigate the effects of different ratios of cardiomyocytes and fibroblasts. Lab Chip. 2021 Oct 26;21(21):4177–95.

17. Xu F, Mao B, Li Y, Zhao Y. Knockdown of HIPK2 attenuates angiotensin II-induced cardiac fibrosis in cardiac fibroblasts. J Cardiovasc Pharmacol. 2022 May 6;

18. Stebler S, Raghunath M. The Scar-in-a-Jar: In Vitro Fibrosis Model for Anti-Fibrotic Drug Testing. Methods Mol Biol Clifton NJ. 2021;2299:147–56.

19. Orozco P, Montoya Y, Bustamante J. Development of endomyocardial fibrosis model using a cell patterning technique: In vitro interaction of cell coculture of 3T3 fibroblasts and RL-14 cardiomyocytes. PLOS ONE. 2020 Feb 24;15(2):e0229158.

20. Wang EY, Smith J, Radisic M. Design and Fabrication of Biological Wires for Cardiac Fibrosis Disease Modeling. Methods Mol Biol Clifton NJ. 2022;2485:175–90.

21. Lee MO, Jung KB, Jo SJ, Hyun SA, Moon KS, Seo JW, et al. Modelling cardiac fibrosis using three-dimensional cardiac microtissues derived from human embryonic stem cells. J Biol Eng. 2019 Dec;13(1):15.

22. Lian X, Hsiao C, Wilson G, Zhu K, Hazeltine LB, Azarin SM, et al. Robust cardiomyocyte differentiation from human pluripotent stem cells via temporal modulation of canonical Wnt signaling. Proc Natl Acad Sci U S A. 2012 Jul 3;109(27):E1848–1857.

23. Allen Institute for Cell Science. Cardiomyocyte Differentiation Methods [Internet]. 2020. Available from: https://www.allencell.org/uploads/8/1/9/9/81996008/sop_for_cardiomyocyte_differentiation_methods_v1.2.pdf

24. Johnson TD, Hill RC, Dzieciatkowska M, Nigam V, Behfar A, Christman KL, et al. Quantification of decellularized human myocardial matrix: A comparison of six patients. Proteomics Clin Appl. 2016 Jan;10(1):75–83.

25. Gwyther TA, Hu JZ, Billiar KL, Rolle MW. Directed cellular self-assembly to fabricate cell-derived tissue rings for biomechanical analysis and tissue engineering. J Vis Exp JoVE. 2011 Nov 25;(57):e3366.

26. Sala L, van Meer BJ, Tertoolen LGJ, Bakkers J, Bellin M, Davis RP, et al. MUSCLEMOTION: A Versatile Open Software Tool to Quantify Cardiomyocyte and Cardiac Muscle Contraction In Vitro and In Vivo. Circ Res. 2018 02;122(3):e5–16.

27. Gwyther TA, Hu JZ, Christakis AG, Skorinko JK, Shaw SM, Billiar KL, et al. Engineered vascular tissue fabricated from aggregated smooth muscle cells. Cells Tissues Organs. 2011;194(1):13–24.

28. Roman B, Kumar SA, Allen SC, Delgado M, Moncayo S, Reyes AM, et al. A Model for Studying the Biomechanical Effects of Varying Ratios of Collagen Types I and III on Cardiomyocytes. Cardiovasc Eng Technol. 2021 Jun 1;12(3):311–24.

29. Lee A, Hudson AR, Shiwarski DJ, Tashman JW, Hinton TJ, Yerneni S, et al. 3D bioprinting of collagen to rebuild components of the human heart. Science. 2019 Aug 2;365(6452):482–7.

30. Rashedi I, Talele N, Wang XH, Hinz B, Radisic M, Keating A. Collagen scaffold enhances the regenerative properties of mesenchymal stromal cells. PLOS ONE. 2017 Oct 31;12(10):e0187348.

31. Kaiser NJ, Kant RJ, Minor AJ, Coulombe KLK. Optimizing Blended Collagen-Fibrin Hydrogels for Cardiac Tissue Engineering with Human iPSC-derived Cardiomyocytes. ACS Biomater Sci Eng. 2019 Feb 11;5(2):887–99.

32. Bott K, Upton Z, Schrobback K, Ehrbar M, Hubbell JA, Lutolf MP, et al. The effect of matrix characteristics on fibroblast proliferation in 3D gels. Biomaterials. 2010 Nov 1;31(32):8454–64.

33. Miron-Mendoza M, Seemann J, Grinnell F. Collagen Fibril Flow and Tissue Translocation Coupled to Fibroblast Migration in 3D Collagen Matrices. Mol Biol Cell. 2008 May;19(5):2051–8.

34. Wilks BT, Evans EB, Howes A, Hopkins CM, Nakhla MN, Williams G, et al. Quantifying Cell-Derived Changes in Collagen Synthesis, Alignment, and Mechanics in a 3D Connective Tissue Model. Adv Sci. 2022;9(10):2103939.

35. Edalat SG, Jang Y, Kim J, Park Y. Collagen Type I Containing Hybrid Hydrogel Enhances Cardiomyocyte Maturation in a 3D Cardiac Model. Polymers. 2019 Apr;11(4):687.

36. Feng Z, Matsumoto T, Nakamura T. Measurements of the mechanical properties of contracted collagen gels populated with rat fibroblasts or cardiomyocytes. J Artif Organs Off J Jpn Soc Artif Organs. 2003;6(3):192–6.

37. Tani H, Kobayashi E, Yagi S, Tanaka K, Kameda-Haga K, Shibata S, et al. Heart-derived collagen promotes maturation of engineered heart tissue. Biomaterials. 2023 Aug 1;299:122174.

38. Karbassi E, Fenix A, Marchiano S, Muraoka N, Nakamura K, Yang X, et al. Cardiomyocyte maturation: advances in knowledge and implications for regenerative medicine. Nat Rev Cardiol. 2020 Feb 3;1–19.

39. González A, López B, Ravassa S, San José G, Díez J. The complex dynamics of myocardial interstitial fibrosis in heart failure. Focus on collagen cross-linking. Biochim Biophys Acta BBA - Mol Cell Res. 2019 Sep 1;1866(9):1421–32.

40. López B, Querejeta R, González A, Larman M, Díez J. Collagen Cross-Linking But Not Collagen Amount Associates With Elevated Filling Pressures in Hypertensive Patients With Stage C Heart Failure. Hypertension. 2012 Sep;60(3):677–83.

41. Jones MG, Andriotis OG, Roberts JJ, Lunn K, Tear VJ, Cao L, et al. Nanoscale dysregulation of collagen structure-function disrupts mechano-homeostasis and mediates pulmonary fibrosis. eLife. 7:xse36354.

