## Supplemental Figures for "Exogenous ECM in an environmentally-mediated *in vitro* model for cardiac fibrosis"

### Slide 1
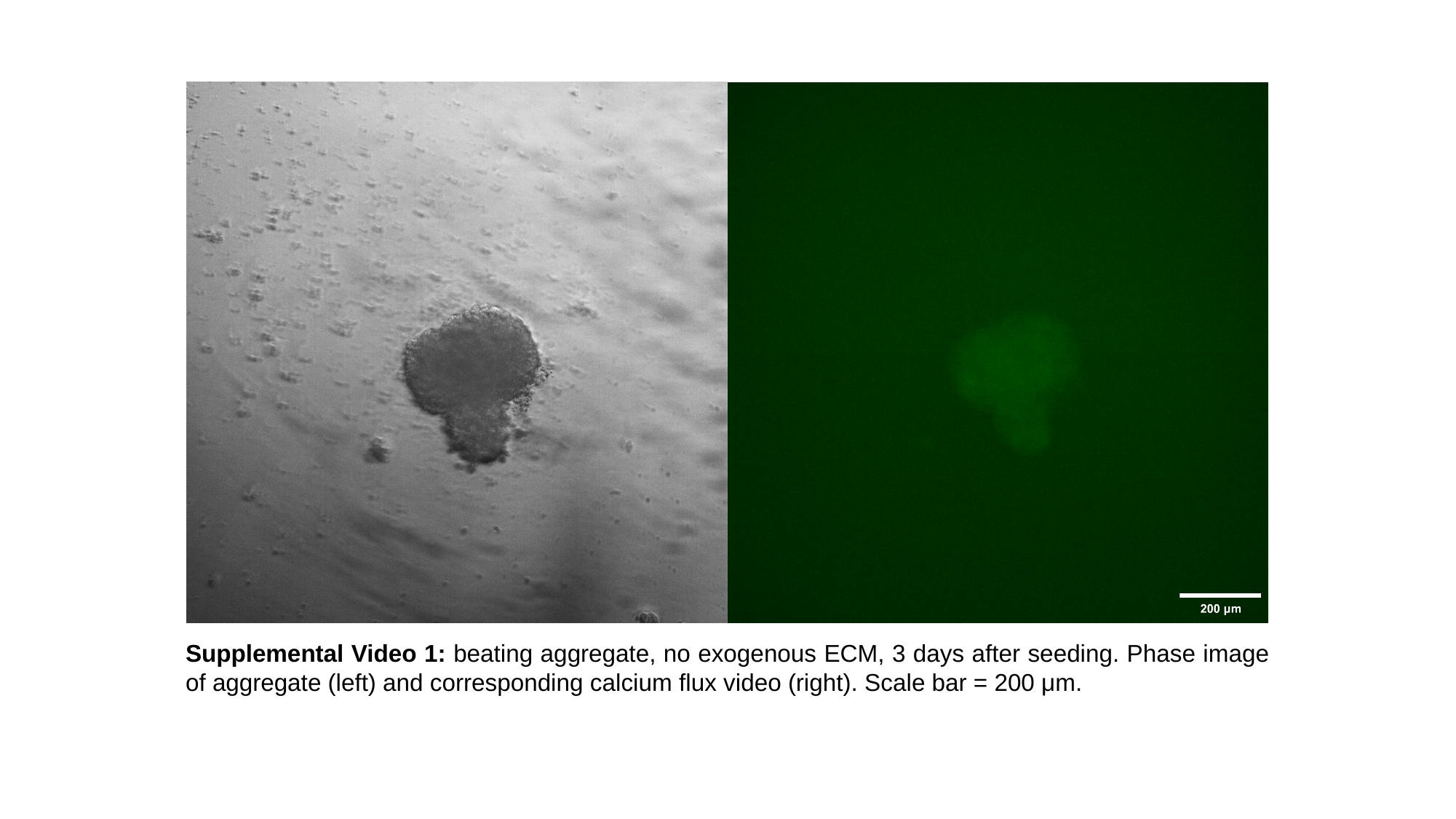

Supplemental Video 1: beating aggregate, no exogenous ECM, 3 days after seeding. Phase image of aggregate (left) and corresponding calcium flux video (right). Scale bar = 200 μm.

### Slide 2
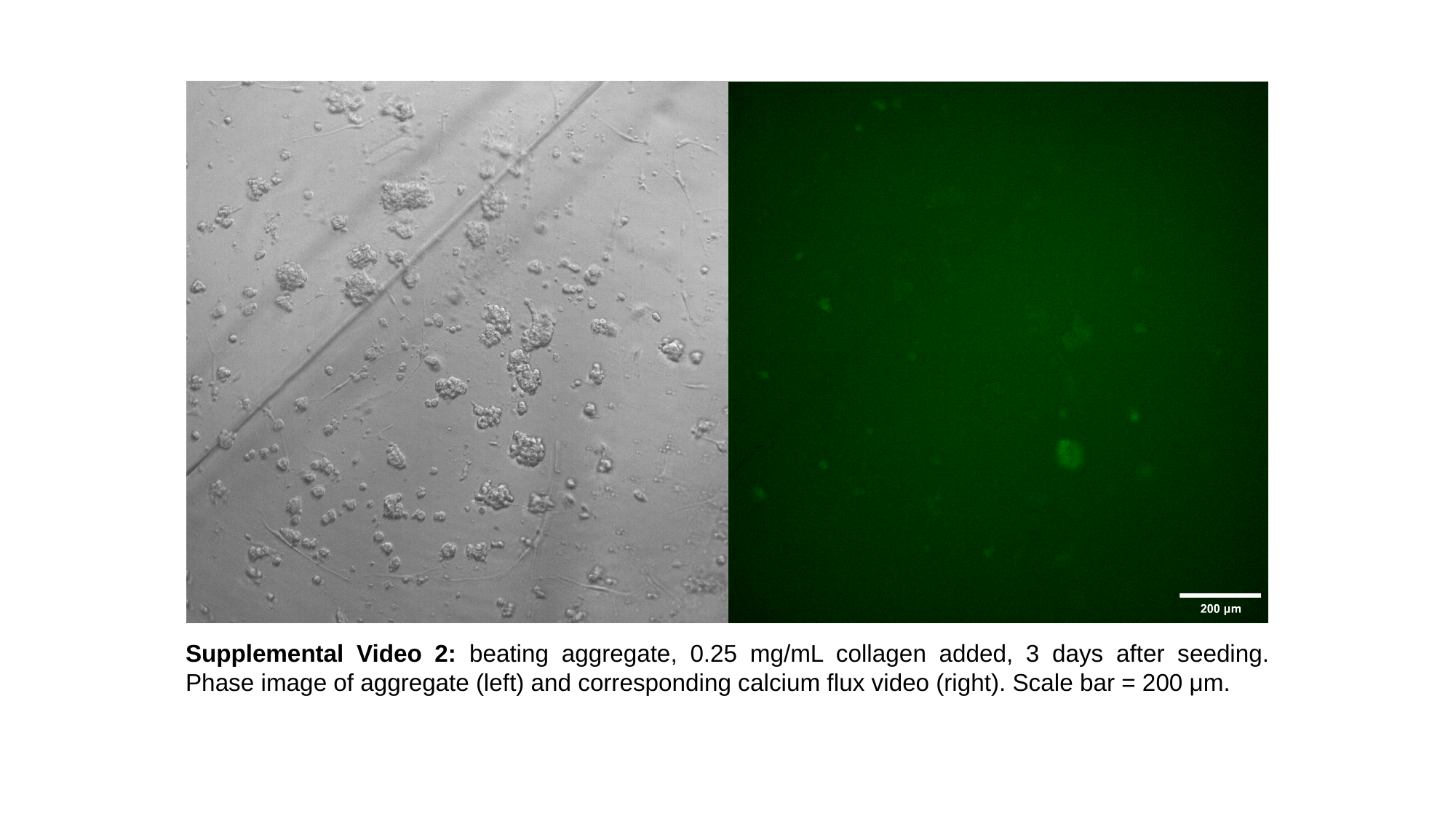

Supplemental Video 2: beating aggregate, 0.25 mg/mL collagen added, 3 days after seeding. Phase image of aggregate (left) and corresponding calcium flux video (right). Scale bar = 200 μm.

### Slide 3
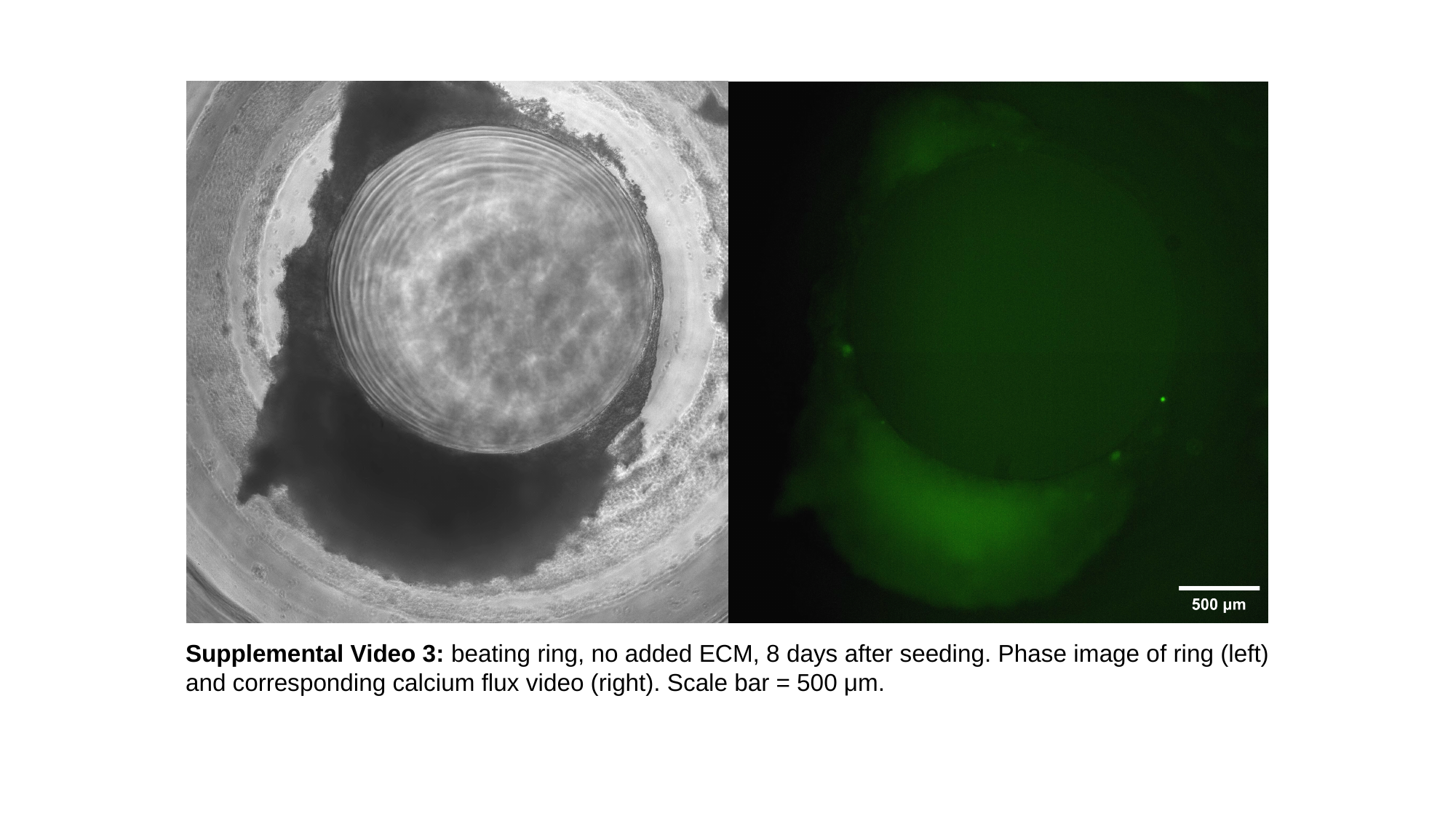

Supplemental Video 3: beating ring, no added ECM, 8 days after seeding. Phase image of ring (left) and corresponding calcium flux video (right). Scale bar = 500 μm.

### Slide 4
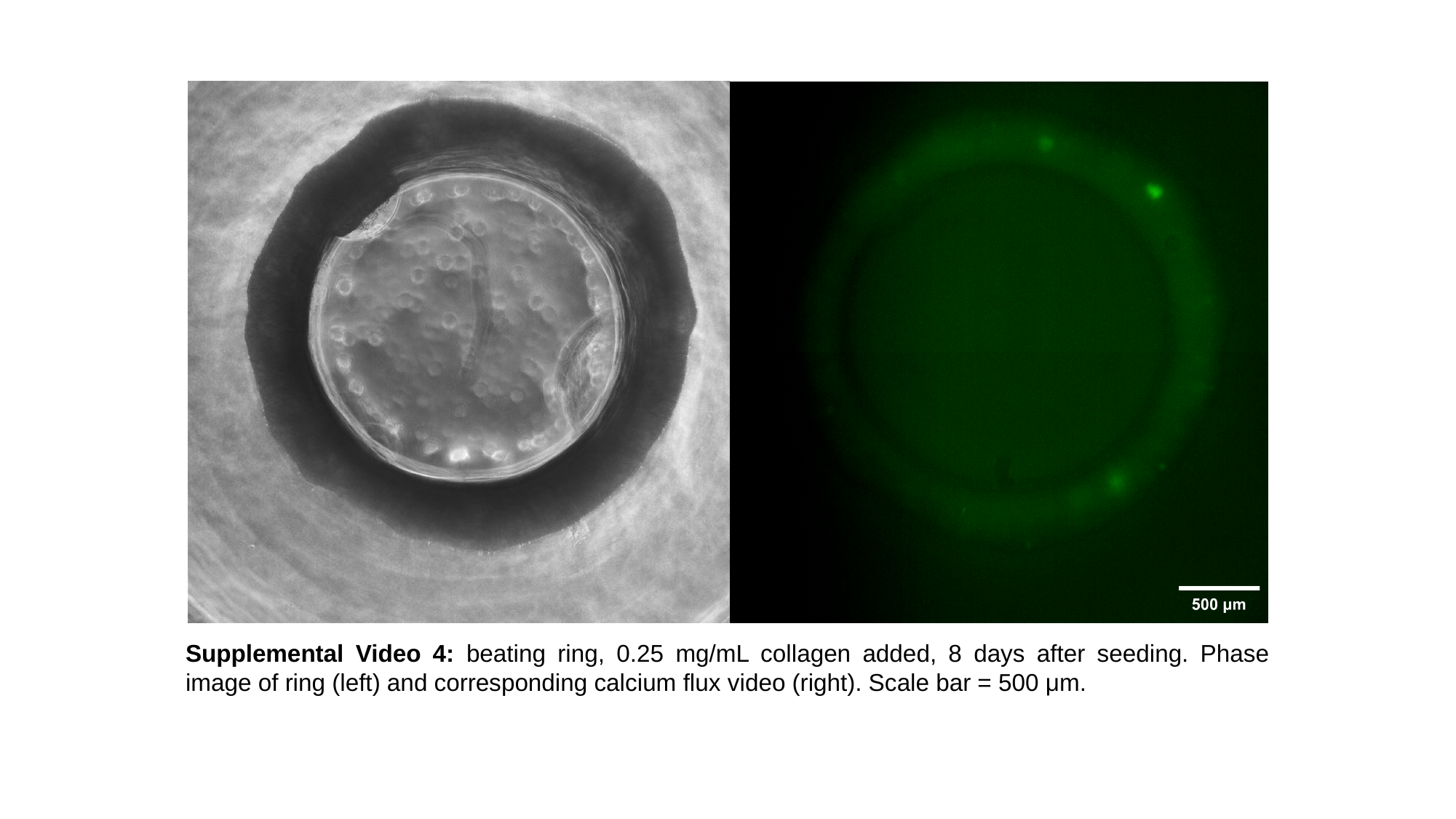

Supplemental Video 4: beating ring, 0.25 mg/mL collagen added, 8 days after seeding. Phase image of ring (left) and corresponding calcium flux video (right). Scale bar = 500 μm.

### Slide 5
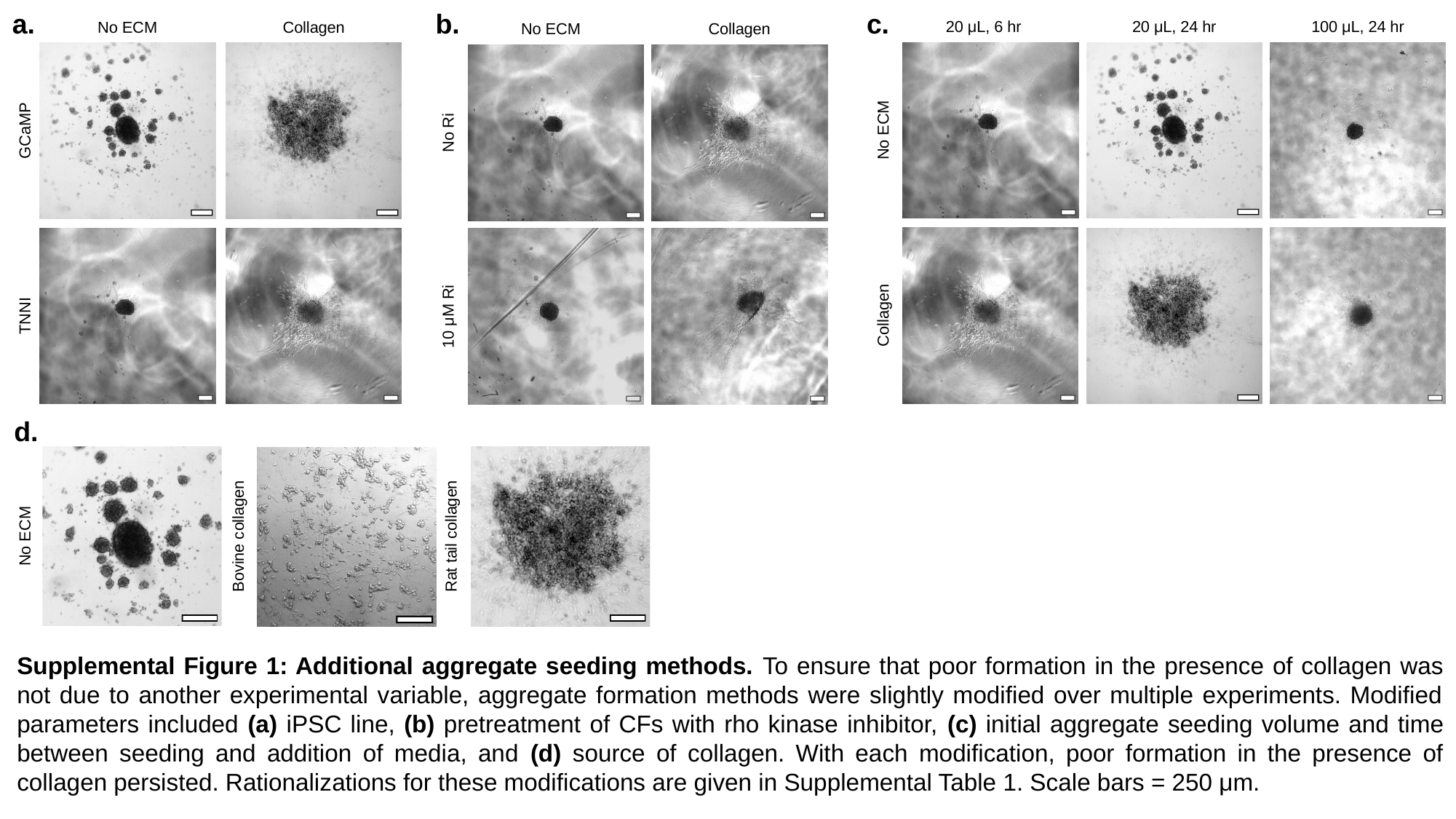

a.
b.
c.
20 μL, 6 hr
20 μL, 24 hr
100 μL, 24 hr
No ECM
Collagen
No ECM
Collagen
GCaMP
TNNI
No ECM
Collagen
No Ri
10 μM Ri
d.
Rat tail collagen
Bovine collagen
No ECM
Supplemental Figure 1: Additional aggregate seeding methods. To ensure that poor formation in the presence of collagen was not due to another experimental variable, aggregate formation methods were slightly modified over multiple experiments. Modified parameters included (a) iPSC line, (b) pretreatment of CFs with rho kinase inhibitor, (c) initial aggregate seeding volume and time between seeding and addition of media, and (d) source of collagen. With each modification, poor formation in the presence of collagen persisted. Rationalizations for these modifications are given in Supplemental Table 1. Scale bars = 250 μm.

### Slide 6
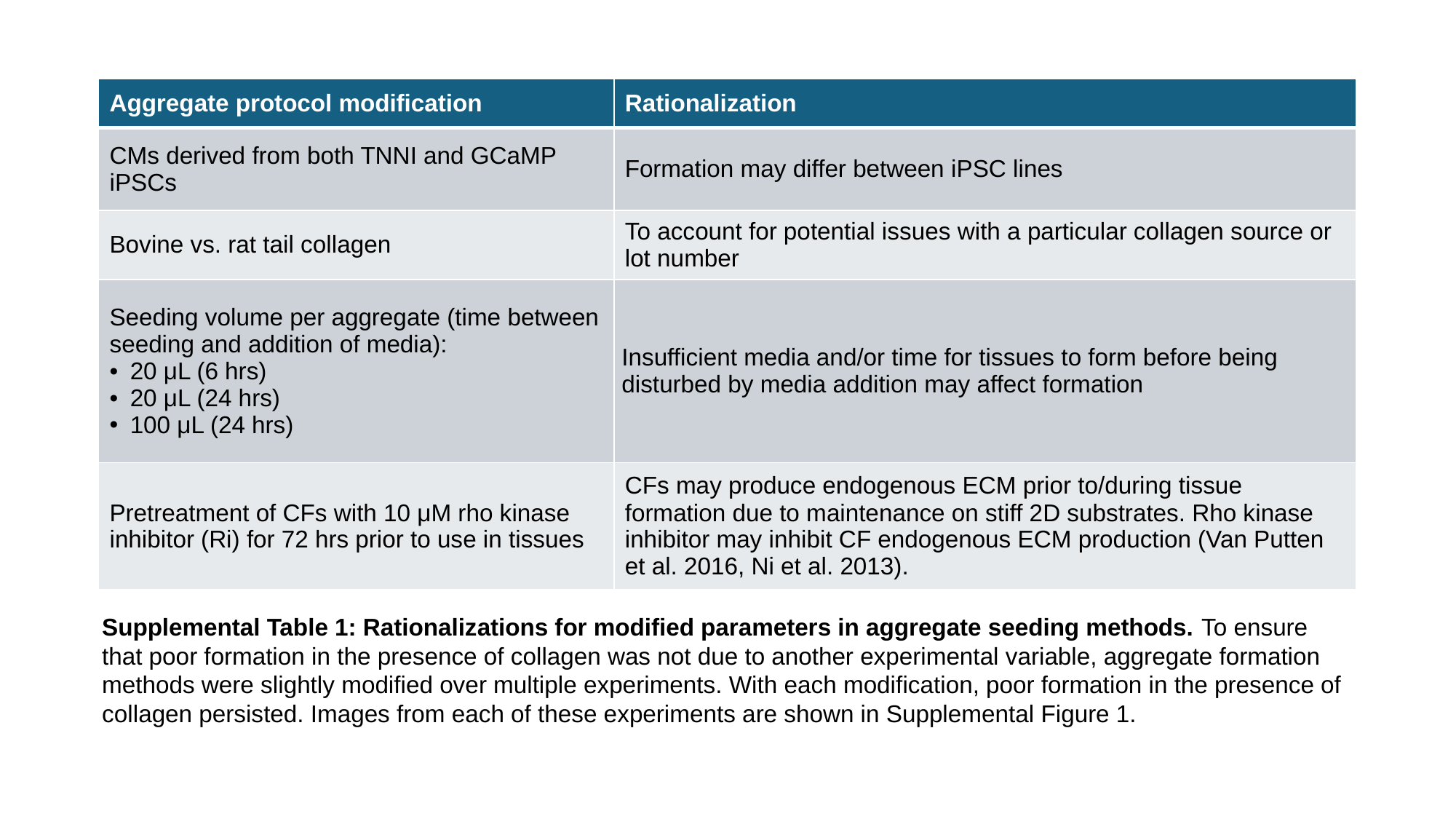

| Aggregate protocol modification | Rationalization |
| --- | --- |
| CMs derived from both TNNI and GCaMP iPSCs | Formation may differ between iPSC lines |
| Bovine vs. rat tail collagen | To account for potential issues with a particular collagen source or lot number |
| Seeding volume per aggregate (time between seeding and addition of media): 20 μL (6 hrs) 20 μL (24 hrs) 100 μL (24 hrs) | Insufficient media and/or time for tissues to form before being disturbed by media addition may affect formation |
| Pretreatment of CFs with 10 μM rho kinase inhibitor (Ri) for 72 hrs prior to use in tissues | CFs may produce endogenous ECM prior to/during tissue formation due to maintenance on stiff 2D substrates. Rho kinase inhibitor may inhibit CF endogenous ECM production (Van Putten et al. 2016, Ni et al. 2013). |
Supplemental Table 1: Rationalizations for modified parameters in aggregate seeding methods. To ensure that poor formation in the presence of collagen was not due to another experimental variable, aggregate formation methods were slightly modified over multiple experiments. With each modification, poor formation in the presence of collagen persisted. Images from each of these experiments are shown in Supplemental Figure 1.

### Slide 7
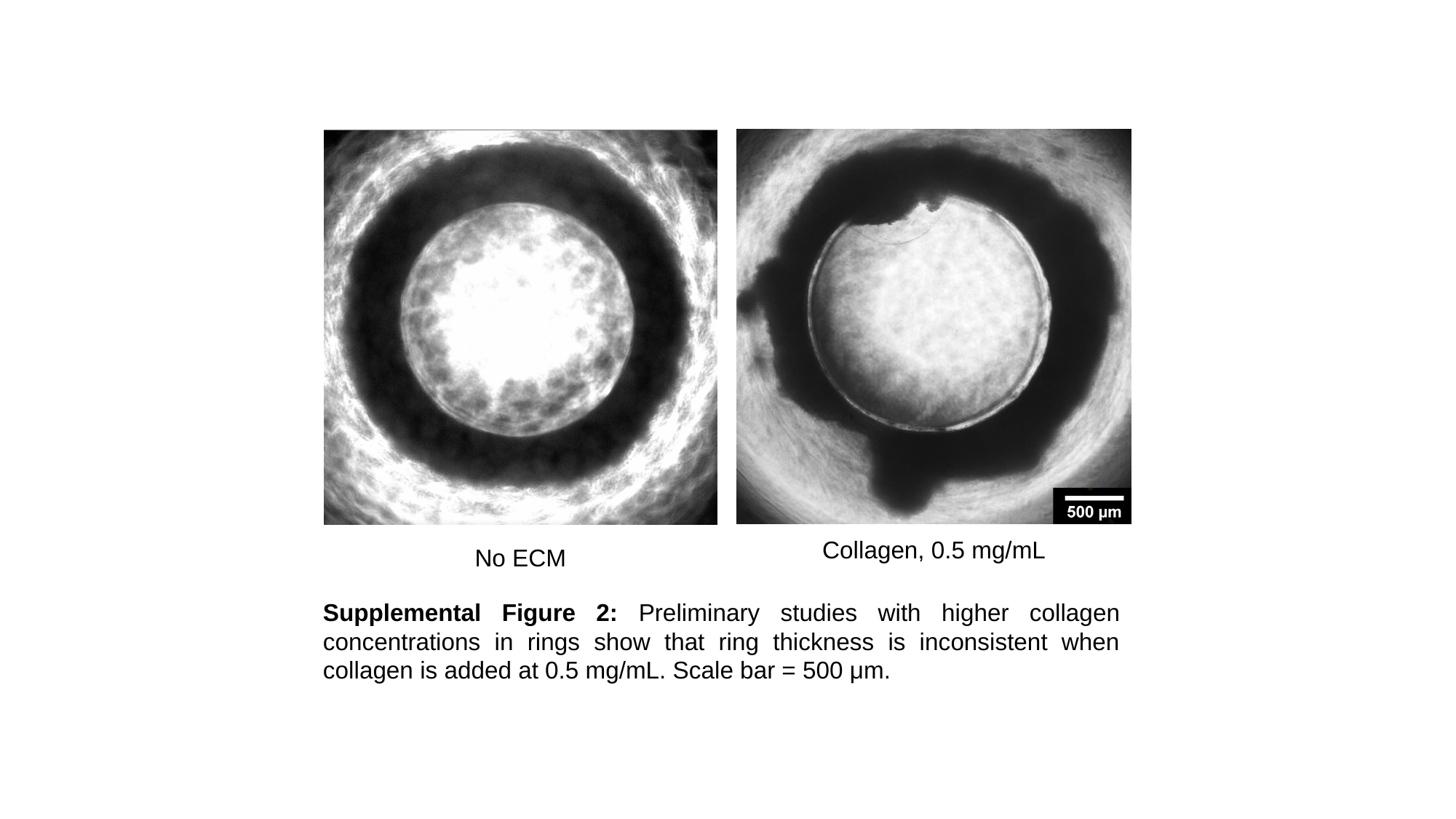

No ECM
Collagen, 0.5 mg/mL
Supplemental Figure 2: Preliminary studies with higher collagen concentrations in rings show that ring thickness is inconsistent when collagen is added at 0.5 mg/mL. Scale bar = 500 μm.

### Slide 8
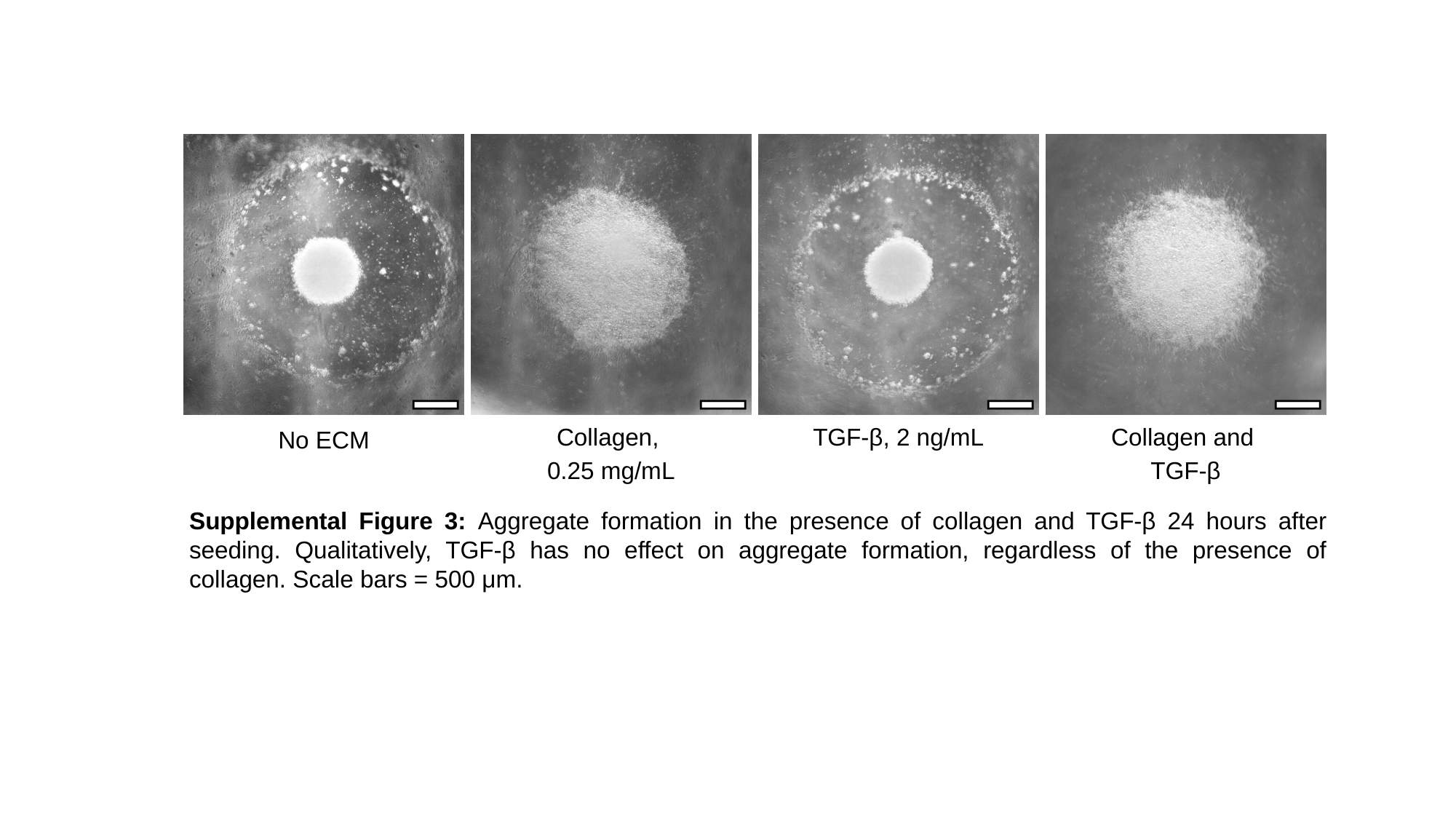

Collagen,
0.25 mg/mL
TGF-β, 2 ng/mL
Collagen and
TGF-β
No ECM
Supplemental Figure 3: Aggregate formation in the presence of collagen and TGF-β 24 hours after seeding. Qualitatively, TGF-β has no effect on aggregate formation, regardless of the presence of collagen. Scale bars = 500 μm.
